# The Non-random Location of Autosomal Genes that Participate in X Inactivation

**DOI:** 10.1101/669929

**Authors:** Barbara R. Migeon

## Abstract

By transcribing XIST RNA, the human inactive X chromosome has a prime role in X-dosage compensation. Yet, the autosomes also play an important role in the process. In fact, multiple genes on human chromosome 1 interact with XIST RNA to silence the inactive Xs, no matter how many there are. And it is likely that multiple genes on human chromosome 19 prevent the silencing of the *single* active X, which is a highly dosage sensitive process. Previous studies of the organization of chromosomes in the nucleus and their genomic interactions indicate that most contacts are intra-chromosomal. Coordinate transcription or dosage regulation could explain the clustered organization of these autosomal genes on these two chromosomes that are critical for X dosage compensation in human cells. Unlike those on chromosome 1, the genes within the critical eight MB region of chromosome 19, have remained together in all mammals assayed, except rodents, indicating that their proximity in non-rodent mammals is evolutionarily conserved.

---

When female mammals compensate for sex differences in the dosage of X linked genes by inactivating X chromosomes, the X chromosome(s) to be silenced has a major role in the process. In all mammals, a non-coding RNA, encoded by the X, is essential to its being inactivated by epigenetic factors [1]. Clearly, the bi-directional spread of Xist RNA from its locus in the middle of the X chromosome initiates the silencing process in eutherian mammals [2, 3]. Once coated with enough Xist RNA, the future inactive X moves toward the nuclear lamina, where its chromatin is transformed from euchromatin to heterochromatin [4, 5]. In addition, the other long non coding RNAs, implicated in the process, that is, the potential Xist repressors, rodent-specific *Tsix*, [6], and the primate specific *XACT* [7], are also encoded by the X chromosome.

The silencing of the future inactive X, or Xs, is attributable to a Rube-Goldberg type of mechanism that not only brings it close to the nuclear periphery (where inactive chromatin tends to reside), but also attracts the epigenetic factors that silence it. Ultimately, the binding of Xist RNA results in expulsion of the SWI/SNF complex from the inactive X [8]. The few active (escape) genes on that X chromosome manage to find their way out of the heterochromatic mass of inactive chromatin towards the center of the nucleus, where transcription occurs [9]. Yet, Xist RNA cannot do this alone, as autosomal gene products are essential to complete the silencing process [4, 5, 10].

In pursuit of autosomal genes that cooperate with the X chromosome, Percec et al., 2003 [11] used ENU chemical mutagenesis to screen for autosomal mutations involved in the initiation of X inactivation in mice. They identified regions of mouse chromosomes 5, 10 and 15, which seemed to affect the choice of the mouse inactive X. More recent studies in mice have elucidated the essential autosomal products that interact with Xist RNA to silence the chromosome [4, 5, 12, 13] (Table 1). These include the lamin B receptor Lbr, the satellite attachment factor A (*Saf-A*) and *Sharp*, (*Smrt* and Hdac Associated Repressor Protein, also called *Spen*). *SPEN, LBR* and *SAFA* map to human chromosome 1; *Lbr* and *Safa* also map to mouse chromosome 1, whereas *Sharp* is on mouse chromosome 4, (orthologous to human chromosome 1). Other genes that have been implicated in the silencing process are *RBM 15* and *SETDB1*, on human chromosome 1, and mouse chromosome 3 – also orthologous to human chromosome 1. Therefore, the genes on human chromosome 1 that play a role in silencing the future inactive X also map to mouse chromosome 1 or its orthologs (*Table 1, Figure 1a*). Conceivably, genes that are on three different chromosomes in mice evolved to be on a single human chromosome to facilitate their interaction in silencing the X.

**TABLE 1:**
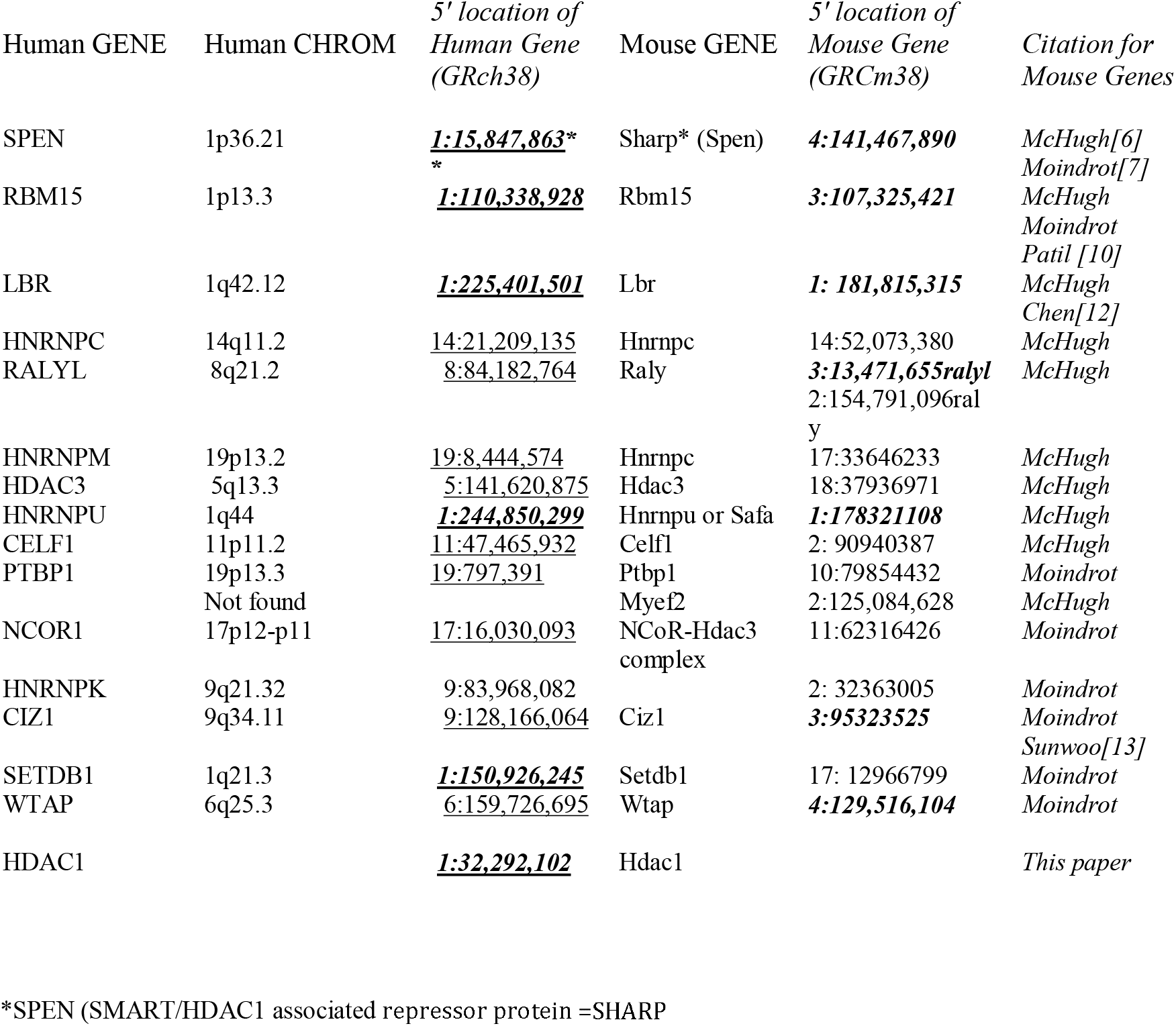
Location of mouse and human genes that silence the inactive X

**Figure 1.**
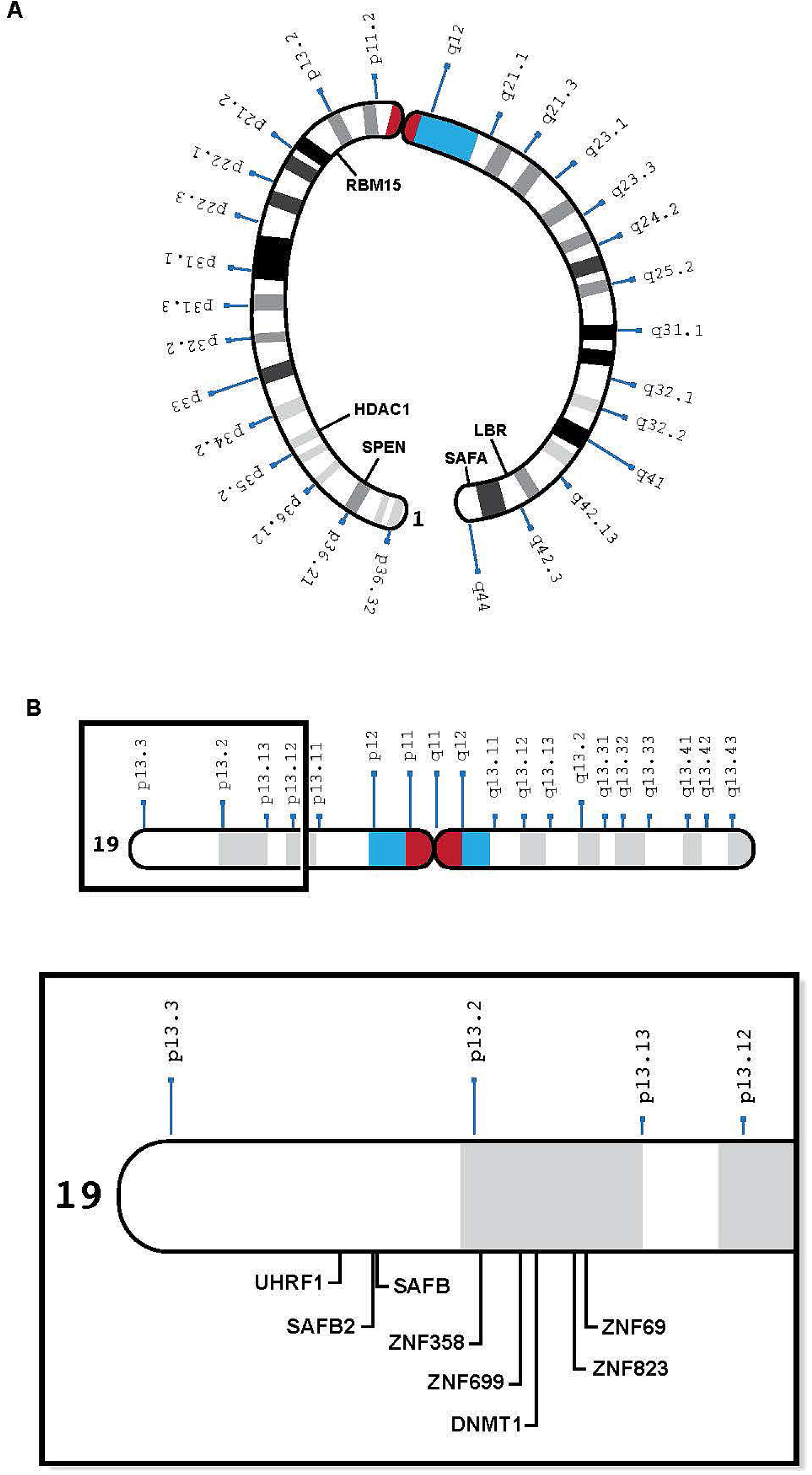
**A**. Human Chromosome 1 with relevant genes, bent to show telomeres in proximity: SPEN 1p36.2, HDAC1 1p35.2 RBM15 1p13.3, LBR 1q42.12, and SAFA, 1q44, *See Table 1*. **B**. Human Chromosome 19 insert with relevant genes, showing proximity of genes in 19p13.1-13.2: UHRF1, Safb2, SAFB, ZNF 358, ZNF 699, SMARCA4, DNMT1, ZNF 823, ZNF 69 *See Table 2*.

In prokaryotes, interactions between genes with a common function are facilitated because such genes are contiguous in the genome, organized into operons, with a common promoter [14]. On the other hand, most eukaryotic genes that interact with each other, do not share promotors, and are less well clustered [15]. Yet, it has become apparent that the spatial arrangement of genes in the mammalian nucleus is non-random; chromosome folding and intermingling enable the proximity of genes that reside on the same chromosome, by looping, and even on different chromosomes, by chromosome clustering. The likely advantage of interactions between genes is coordination of their expression – perhaps in the same transcription factory, thought to occur in a discrete nuclear region [16].

Based on HI-C studies of the human genome [17], Thevenin et al. [18] showed that a significant number of functional groups (pairs of interacting proteins, complexes and pathways) are either located within the same chromosome or dispersed over several chromosomes. Those on different chromosomes tend to co-localize in space. In order to coordinate their expression, genes that function together tend to reside on fewer chromosomes than expected by chance. On the same chromosome, they are closer to each other than randomly chosen genes; on different chromosomes, they tend to be closer to each other in 3D space [18]. Among the best documented inter-chromosomal interactions are those between the mouse X chromosomal gene, *Xist*, and the autosomal epigenetic factors mentioned above that help silence the X chromosome from which the up-regulated *Xist* locus is being transcribed [15].

When extending her observations in mice to other mammals, Lyon suggested there was only a single *active* X, no matter the number of X’s in a cell [19]; however, the literature has persisted in labeling the mammalian process of X dosage compensation, *X inactivation*, which focuses us on the process of silencing the inactive X. Therefore, the salient question has been, “How does one *choose* the X chromosome that becomes *inactive?*” Because Xist RNA is able to silence any chromosome into which it is inserted [20, 21], it is surprising that few ask the pertinent question, “What protects the single active X from silencing by its own *Xist* locus?”[22, 23].

Further, it has not been easy to show how the mouse *inactive* X is chosen. Earlier studies suggested that an infrequent physical association (kissing) between the *Xist* loci of the two X chromosomes in mouse embryos determined the choice of inactive X [24, 25], but more recent studies indicate that neither the expression of *Xist* nor *Tsix*, its antisense RNA, is affected by the interaction [26, 27].

In addition, Inoue et al. [28] and Harris et al. [29] recently showed that in mice, the choice of *active* X is determined prenatally. Having been imprinted during oocyte differentiation [28, 29], (as predicted by Lyon and Rastan [30]), the active X is always *maternal* in trophectoderm – the first tissue to undergo dosage compensation in the mouse embryo. Because X inactivation in the placenta occurs relatively early in mice, it is likely that the paternal X hasn’t had time to erase the inactivation imprint imposed during the early stages of spermatogenesis [31]. It remains to be seen if the rodent specific Tsix RNA, which is transcribed only from the maternal X in trophectoderm, protects the active X, regardless of its parental origin, from silencing by *Xist* in other mouse embryonic tissues.

Because human oocytes do not express *PCR2*, which imprints the mouse oocyte, [29] and the human maternal X is not imprinted [32], and because human *TSIX* is ineffective, having been truncated during human evolution [32], another means of repressing the *XIST* locus on the future *active* human X is needed to protect it from being silenced. Therefore, to repress its *XIST* locus, recent studies suggest that the future human *active* X needs to interact with human chromosome 19. [23]. These studies reveal a previously unsuspected eight MB region on the short arm of human chromosome 19 (19p13.3-13.2) which contains at least one dosage sensitive gene that is likely to play a role in silencing the *XIST* locus on *one* X chromosome in each cell, to protect it from heterochromatization [22, 23], (Table 2). Candidate genes include satellite attachment factors *SAFB* and *SAFB2*, a cluster of zinc finger proteins that surround *DNMT1* and its co-factor *UHRF1*, among many others. Although most of the zinc finger proteins clustered in the relevant region of human chromosome 19 arose after the split between rodents and humans, the other genes in this region can be found on mouse chromosomes 8,9 and 17, which are orthologous to human chromosome 19 (*Table 2, Figure 1b*). Again, perhaps human 19 evolved to facilitate the interaction of genes that protect the future active X.

**TABLE 2:** Location of mouse and human genes that maintain the active X

| Human GENE | Human CHROM | 5' location of Human Gene (GRch38 ) | Mouse GENE | 5' location of Mouse Gene (GRCm38) |
| --- | --- | --- | --- | --- |
| SAFB | 19p13.3 | ***19: 5,623,034\*\**** | Safb | ***17:56, 584,830*** |
| SAFB2 | 19p13.3 | ***19: 5,586,992*** | Safb2 | ***17:56, 560,965*** |
| DNMT1 | 19p13.2 | ***19:10,133,343*** | Dnmt1 | ***9:20,907,209*** |
| UHRF1 | 19p13.3 | ***19: 4,903,079*** | Uhrf1 | ***17: 56, 303,367*** |
| HNRNPM | 19p13.2 | ***19: 8,444,574*** | Hnrnpm | ***17: 33, 646,233*** |
| MBD3 | 19p13.3 | ***19: 1,576,670*** | Mbd3 | 10: 80,392,539 |
| MBD3L-5L | 19p13.2 | ***19: 8,842,392*** | Mbd3l | ***9: 18,478, 359*** |
| PRMT4 or CARM1 | 19p13.2 | ***19:10,871,576*** | Carm1 | ***9: 21,546,894*** |
| ZNF69 | 19p13.2 | ***19:11,887,772*** | Not found |  |
| ZNF823 | 19p13.2 | ***19:11,832,080*** | " |  |
| ZNF443 | 19.p13.2 | ***19:12,4*** | Znf 709 | 8:71,882,068 |
| ZNF699 | 19p13.2 | ***19: 9,291,139*** | Not found |  |
| ZNF627 | 19p13.2 | ***19:11,575,254*** | Znf 867 | 11:59, 461,197 |
| ZNF44 | 19p13.2 | ***19:12,224,685*** | Not found |  |
| ZNF358 pli.87 | 19p13.2 | ***19: 7,580,178*** | Zfp358 | 8: 3,493,138 |

\*\* ***Bold italics***: Human chromosome 19 or mouse orthologs of human chromosome 19

In the genomics era, geneticists tend not to think about the individual autosomes which encode genes of interest; therefore, I was surprised to see that many of the major players that interact with *XIST* to silence the X are encoded by human chromosome 1 [23], *Table 1 & Figure 1a*)], and in the mouse, by the three orthologs of chromosome 1 (chromosomes 1, 3 and 4) (Table 1). In mice, these genes are bound to *Xist* at the same developmental stage [4]. To my knowledge, no one has examined the *Xist*-autosomal interactions by RNA FISH to determine if there is clustering of the three murine chromosome 1 orthologs. The positions of these genes on human chromosome 1 is of interest as some of the genes are present on opposing ends of the chromosome, which would require a large fold in the chromosome to facilitate any interaction (*Figure 1a*). Such intermingling and folding are frequently observed in the 3D nuclear space [17].

Table 3 presents conservation data obtained from the UCSC Genome Browser; it shows that of four relevant genes on chromosome 1, only SAFA and LBR have been on the chromosome since we evolved from marsupials. SPEN and RBM15 although on the same chromosome as SAFA and LBR in primates, are on other chromosomes in marmosets and non-primate mammals. In contrast, except in rodents (rat, mouse and rabbit), the region on chromosome 19 is preserved in primates such as gorilla, orangutang, and marmoset, and other mammals such as cat, dog, pig, horse, cow, and opposum, (Table 4). The exceptional genes include the long noncoding RNA, TINCR, and the MD3L3-5, methyl CPG binding domain proteins, which are on chromosome 19 in primates and in marmoset but are not found in all mammals. The conserved cluster in pig, horse and cow is in the reverse orientation (Table 4). These differences interrupt what would otherwise be an exceptionally long synteny block, but the preservation of so many genes in this region, in spite of multiple evolutionary structural alterations, suggests that the local landscape may be important to function. That the chromosome 19 genes in rodents are not conserved as a group argues that their process of ensuring that one X will remain active differs from that of other mammals [33], perhaps because only rodents have *Tsix* to protect the active X from silencing by *Xist*.

**TABLE 3:**
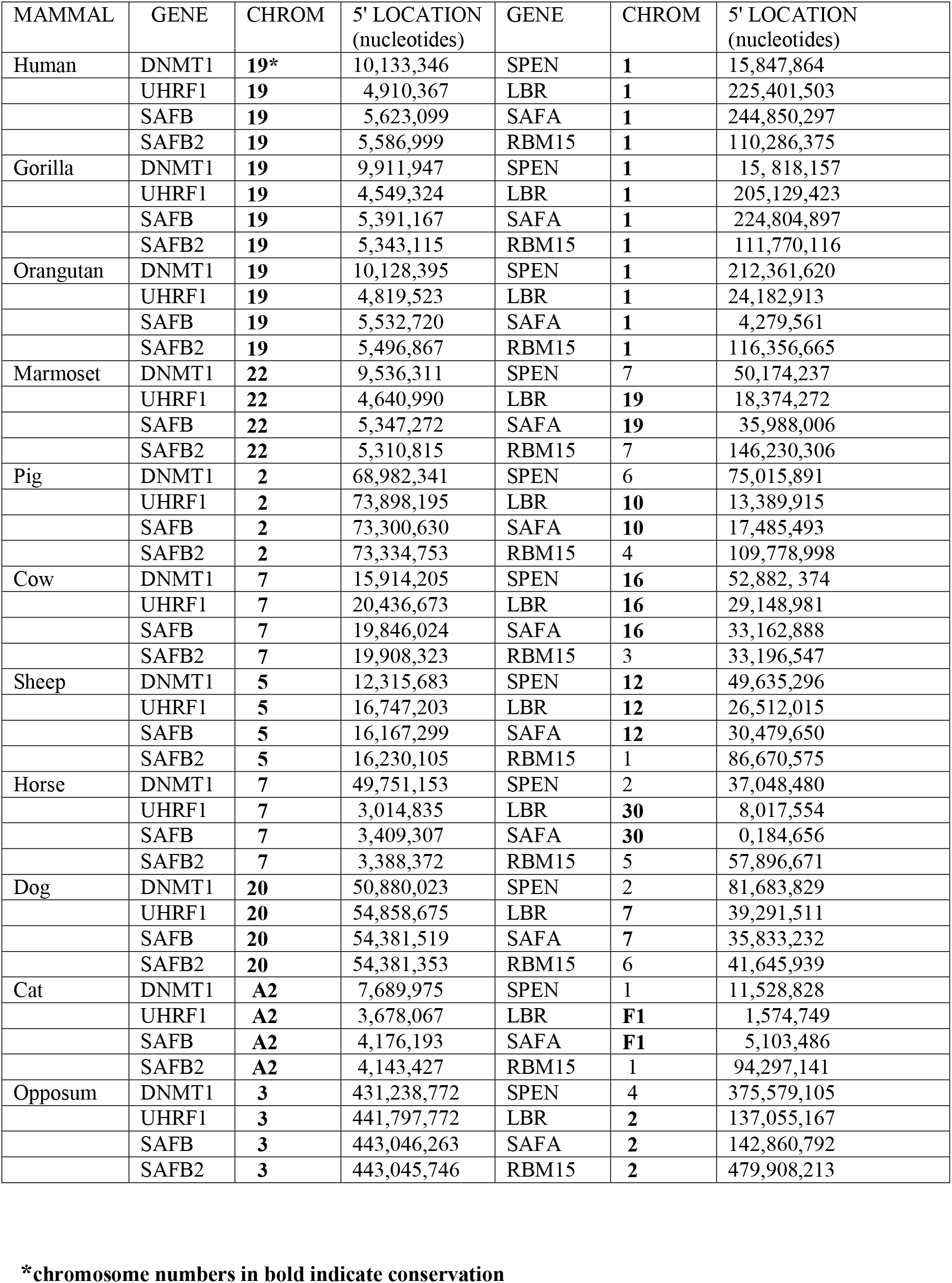
Conservation of Some Candidate Genes, and Not Others in Various Mammals

**TABLE 4:** Site of 18 Clustered Human Chromosome 19 Genes in Other Mammals

| MAMMAL | GENE | CHROMOSOME | SITE 5' (nucleotide) | Other Site(s) |
| --- | --- | --- | --- | --- |
| HUMAN | SIRT6 | 19 | 4,174,109 |  |
|  | PLIN3 | 19 | 4,852,208 |  |
|  | UHRF1 | 19 | 4,910,367 |  |
|  | KDM4B | 19 | 4,969,121 |  |
|  | TINCR | 19 | 5,560,774 |  |
|  | RFX2 | 19 | 5,993,164 |  |
|  | VAV1 | 19 | 6,772,726 |  |
|  | MBD3L4 | 19 | 7,037,748 |  |
|  | INSR | 19 | 7,112,226 |  |
|  | ZNF358 | 19 | 7,516,118 |  |
|  | MAP2K7 | 19 | 7,903,891 |  |
|  | FBN3 | 19 | 8,130,286 |  |
|  | HNRNPM | 19 | 8,269,278 |  |
|  | ZNF558 | 19 | 8,806,170 |  |
|  | OLFM2 | 19 | 9,853,718 |  |
|  | DNMT1 | 19 | 10,133,346 |  |
|  | DNM2 | 19 | 10,828,755 |  |
|  | CARM1 | 19 | 10,871,513 |  |
| ORANGUTAN | SIRT6 | 19 | 4,083,376 |  |
|  | PLIN3 | 19 | 4,752,733 |  |
|  | UHRF1 | 19 | 4,819,523 |  |
|  | KDM4B | 19 | 4,940,648 | also 11 |
|  | TINCR | 19 | 5,468,562 |  |
|  | RFX2 | 19 | 5,907,338 |  |
|  | VAV1 | 19 | 6,738,253 |  |
|  | MBD3L4 | 19 | 7,005,357 |  |
|  | INSR | 19 | 7,065,165 |  |
|  | ZNF358 | 19 | 7,328,128 |  |
|  | MAP2K7 | 19 | 7,862,957 |  |
|  | FBN3 | 19 | 8,037,199 | also 5 |
|  | HNRNPM | 19 | 8, 412,645 |  |
|  | ZNF558 | 19 | 8,801,446 |  |
|  | OLFM2 | 19 | 9,841,684 |  |
|  | DNMT1 | 19 | 10,128,395 |  |
|  | DNM2 | 19 | 10,719,521 |  |
|  | CARM1 | 19 | 10,872.517 |  |
| MARMOSET | SIRT6 | 22 | 3,843,381 |  |
|  | PLIN3 | 22 | 4,576,676 |  |
|  | UHRF1 | 22 | 4,640,990 |  |
|  | KDM4B | 22 | 4,753,547 |  |
|  | TINCR | 22 | 5,280,800 | also 22:19,138,083 |
|  | RFX2 | 22 | 5,714,269 | also 22:13,197,841 and 1:104,992,215 |
|  | VAV1 | 22 | 6,482,055 |  |
|  | MBD3L4 | 22 | 6,745,638 |  |
|  | INSR | 22 | 6,884,705 |  |
|  | ZNF358 | 22 | 7,258,135 | also 16 |
|  | MAP2K7 | 22 | 7,564,197 |  |
|  | FBN3 | 22 | 7,702,224 | also 10 & 2 |
|  | HNRNPM | 22 | 8,116,508 |  |
|  | ZNF558 | 22 | 8,418,995 | also 2 |
|  | OLFM2 | 22 | 9,242,165 | also 5 |
|  | DMNT1 | 22 | 9,536,311 |  |
|  | DNM2 | 22 | 10,141,800 |  |
|  | CARM1 | 22 | 10,298,967 |  |
| PIG | SIRT6 | 2 | 74,568,548 |  |
|  | PLIN3 | 2 | 73,970,200 |  |
|  | UHRF1 | 2 | 73,898,195 |  |
|  | KDM4B | 2 | 73,747,610 |  |
|  | RFX2 | 2 | 72,949,979 |  |
|  | *TINCR* | *not found* |  |  |
|  | VAV1 | 2 | 72,327,498 |  |
|  | MBD3L4 | 2 | 72,012,690 |  |
|  | INSR | 2 | 71,797,542 |  |
|  | ZNF358 | 2 | 71,615,476 |  |
|  | MAP2K7 | 2 | 71,298,318 |  |
|  | FBN3 | 2 | 71,104,118 |  |
|  | HNRNPM | 2 | 70,813,749 |  |
|  | ZNF558 | 2 | (70,582,106) |  |
|  | OLFM2 | 2 | 68,734,136 |  |
|  | DMNT1 | 2 | 68,982,341 |  |
|  | DNM2 | 2 | 69,474,069 |  |
|  | CARM1 | 2 | 69,602,214 |  |
| HORSE | SIRT6 | 7 | 2,539,099 | also 21 |
|  | PLIN3 | 7 | 2,972,664 |  |
|  | UHRF1 | 7 | 3,014,835 |  |
|  | KDM4B | 7 | 3,087,218 | also 23 & 2 |
|  | RFX2 | 7 | 3,649,694 | also 23 |
|  | *TINCR* | *not found* |  |  |
|  | VAV1 | 7 | 4,329,609 |  |
|  | *MBD3L4* | 7 | 52,446,746 | also X |
|  | INSR | 7 | 4,882,687 |  |
|  | ZNF358 | 7 | 4,701,725 | also 4,10, &13 |
|  | MAP2K7 | 7 | 5,229,948 |  |
|  | FBN3 | 7 | 5,361,278 | also 1 & 14 |
|  | HNRNPM | 7 | 52,895,099 |  |
|  | *ZNF558* | 7 | 52,539,233yes |  |
|  | OLFM2 | 7 | 49,967,570yes | also 25 |
|  | DMNT1 | 7 | 49,751,153 |  |
|  | DNM2 | 7 | 49,316,987 | also 25 |
|  | CARM1 | 7 | 49,257,318 |  |
| COW | SIRT6 | 7 | 21,079,141 |  |
|  | PLIN3 | 7 | 20,507,000 |  |
|  | UHRF1 | 7 | 20,436,673 |  |
|  | KDM4B | 7 | 20,308,693 | also 3,8 & 15 |
|  | RFX2 | 7 | 19,126,799 |  |
|  | *TINCR* | *not found* |  |  |
|  | VAV1 | 7 | 18,866,379 | also 3 & 11 |
|  | MD3L4 | 7 | 17,264,390 | also 11 |
|  | INSR | 7 | 17,276,143 | also 21 & 3 |
|  | ZNF358 | 7 | 17,610,070 | also 4,14,17 & 18 |
|  | MAP2K7 | 7 | 17,891,887 |  |
|  | FBN3 | 7 | 18,005,675 | also 10 & 7:26,698,333 |
|  | HNRNPM | 7 | 18,289,395 | also 11 |
|  | ZNF558 | 7 | 17,220,537 | also 19,11 & 25 |
|  | OLFM2 | 7 | 15, 550,353 |  |
|  | DNMT1 | 7 | 15,914,205 |  |
|  | DNM2 | 7 | 16,465,942 | also 11 |
|  | CARM1 | 7 | 16,571,428 |  |
| DOG | SIRT6 | 20 | 55,416,563 |  |
|  | PLIN3 | 20 | 54,924, 119 |  |
|  | UHRF1 | 20 | 54,858,675 |  |
|  | KDM4B | 20 | 54, 715,308 | also 11, 15, & 21 |
|  | RFX2 | 20 | 54,013, 618 | also 1 |
|  | *TINCR* | *not found* |  |  |
|  | VAV1 | 20 | 53,482, 255 |  |
|  | MBD3L4 | 20 | 53,213,540 |  |
|  | INSR | 20 | 52,017, 347 |  |
|  | ZNF358 | 20 | 52,314,421 |  |
|  | MAP2K7 | 20 | 52,594, 536 |  |
|  | FBN3 | 20 | 52,723,997 | also 11 & 30 |
|  | HNRNPM | 20 | 52,997,963 |  |
|  | ZNF558 | 20 | 51, 897, 297 |  |
|  | OLFM2 | 20 | 51,148, 154 |  |
|  | DMNT1 | 20 | 50,880,023 |  |
|  | DNM2 | 20 | 50,399,784 |  |
|  | CARM1 | 20 | 50, 331,081 |  |
| CAT | SIRT6 | A2 | 3,162,759 |  |
|  | PLIN3 | A2 | 3,631,793 |  |
|  | UHRF1 | A2 | 3,678,067 |  |
|  | KDM4B | A2 | 3,765,143 |  |
|  | RFX2 | A2 | 4,427,650 |  |
|  | *TINCR* | *not found* |  |  |
|  | VAV1 | A2 | 5,108,402 |  |
|  | *MBD3L4* | A2 | 5,395,765 |  |
|  | INSR | A2 | 6,443,171 |  |
|  | ZNF358 | A2 | 6,267,306 |  |
|  | MAP2K7 | A2 | 6,004,415 |  |
|  | FBN3 | A2 | 5,820,368 | also A1 |
|  | HNRNPM | A2 | 5,569,560 |  |
|  | ZNF558 | A2 | 6,657,696 |  |
|  | OLFM2 | A2 | 7,484,626 |  |
|  | DMNT1 | A2 | 7,689,975 |  |
|  | DNM2 | A2 | 8,118,334 |  |
|  | CARM1 | A2 | 8,257,736 |  |
| OPOSSUM | SIRT6 | 3 | 440,652,009 |  |
|  | PLIN3 | 3 | 441,702,797 |  |
|  | UHRF1 | 3 | 441,797,772 |  |
|  | KDM4B | 3 | 441,910,670 |  |
|  | RFX2 | 3 | 443,674,276 |  |
|  | *TINCR* | *not found* |  |  |
|  | VAV1 | 3 | 444,980,624 |  |
|  | *MBD3L4* | *not found* |  |  |
|  | INSR | 3 | 463,520,164 |  |
|  | ZNF388 | *unassigned* |  |  |
|  | MAP2K7 | 3 | 462,757,443 |  |
|  | FBN3 | **3** | 461,508,720 |  |
|  | HNRNPM | 3 | 460,359,655 |  |
|  | *ZNF558* | **4** | 409,014,310 |  |
|  | OLFM2 | 3 | 431,554,923 |  |
|  | DMNT1 | 3 | 431,238,772 |  |
|  | DNM2 | 3 | 430,280, 994 |  |
|  | CARM1 | 3 | 430,212,862 |  |
| MOUSE | SIRT6 | 10 | 81,621,787 |  |
|  | PLIN3 | 17 | 56,277,475 |  |
|  | UHRF1 | 17 | 56,304,407 |  |
|  | KDM4B | 17 | 56,326,074 |  |
|  | RFX2 | 17 | 56,775,897 |  |
|  | *TINCR* | 17 | 56,551,534 |  |
|  | VAV1 | 17 | 57,279,100 |  |
|  | *MBD3L4* | *not found* |  |  |
|  | INSR | 8 | 3,150,922 |  |
|  | ZNF358 | 8 | 3,493,154 |  |
|  | MAP2K7 | 8 | 4,238,740 |  |
|  | FBN3 | 18 | 58,012,265 |  |
|  | HNRNPM | 17 | 33,646,236 |  |
|  | *ZNF558* | *not found* |  |  |
|  | OLFM2 | 9 | 20,672,332 |  |
|  | DNMT1 | 9 | 20,,907,209 |  |
|  | DNM2 | 9 | 21,425,244 |  |
|  | CARM1 | 9 | 21,546,894 |  |
| RAT | SIRT6 | 7 | 10,937,622 |  |
|  | PLIN3 | 9 | 10,774,869 |  |
|  | UHRF1 | 9 | 10,738,211 |  |
|  | KDM4B | 9 | 10,656,035 |  |
|  | RFX2 | 9 | 10,216,249 |  |
|  | *TINCR* | 9 | 10,499,290 |  |
|  | VAV1 | 9 | 9,617,783 |  |
|  | *MBD3L4* | 8 | 18,226,238 | 2014 assembly |
|  | INSR | 12 | 1,678,623 |  |
|  | ZNF358 | 12 | 2,046,542 |  |
|  | MAP2K7 | 12 | 2,546,139 |  |
|  | FBN3 | 18 | 53,070,463 |  |
|  | HNRNPM | 7 | 18,516,253 |  |
|  | ZNF558 | *not found* |  |  |
|  | OLFM2 | 8 | 21,684,494 |  |
|  | DNMT1 | 8 | 21,922,515 |  |
|  | DNM2 | 8 | 22,458,869 |  |
|  | CARM1 | 8 | 22,527,213 |  |
| RABBIT | SIRT6 | 3 | 16,044,566 |  |
|  | PLIN3 | *not found* |  |  |
|  | UHRF1 | 1 | 47,672,908 |  |
|  | KDM4B | 1 | 47,085,460 | Also 13 & 12 &<br>1:121,843,123 |
|  | RFX2 | 1 | 51,045,589 |  |
|  | TINCR | *not found* |  |  |
|  | VAV1 | 13 | 56,144,807 |  |
|  | MBD3L4 | *unknown* |  |  |
|  | INSR | un0069 | 1,077,773 |  |
|  | ZNF358 | un0069 | 914,737 |  |
|  | MAP2K7 | un0069 | 665,019 |  |
|  | FBN3 | un0069 | 502,497 | Also 3:11,898,428 |
|  | HNRNPM | un0069 | 252,960 | Also X |
|  | ZNF558 | *not found* |  |  |
|  | OLFM2 | un0135 | 375,668 |  |
|  | DNMT1 | un0135 | 156,550 |  |
|  | DNM2 | 13 | 20,368,794 |  |
|  | CARM1 | 1 | 51,421,465 |  |

Most likely, the relevant genes on the same chromosome are co-regulated. The advantage of genes clustered in interphase is that they can be programmed for simultaneous transcription. To silence *XIST* on the future active X, some genes in the chromosome 19 cluster might be transcribed together, perhaps if they are close enough in 3D space, as a single transcript. The telomeric location of genes on primate chromosome 1 that participate in *XIST* silencing (*Figure 1a*) suggest that the two ends of the chromosome might physically interact at the time of transcription.

Several important questions remain unanswered: First, how do multiple genes in the inactivation (activation) pathway on human chromosome 1 (or chromosome 19) coordinately interact with each other? And then, how do autosomal genes encoding protein products, interact with the X chromosome?

Recent studies suggest that the intra-chromosomal gene interactions occur within the same topologically-associating-domain (TAD)[34, 35] and that TADS align with co-coordinately regulated gene clusters, fostering long-range contacts and preventing deleterious interactions between genes in different TADs [35] One would like to examine the candidate genes on human chromosomes 1 and 19, at the appropriate time in development, to determine if they are located within the same TAD, or are otherwise coordinately regulated. It is unlikely that the occurrence of multiple silencers of the inactive X on human chromosome 1 and *XIST* repressors on human chromosome 19 is coincidental.

The question of how genes on an autosome interact with the genes on the X chromosome is especially challenging because in the human species either one or several X chromosomes can be silenced within a cell, the number dependent upon the number of X chromosomes in the genome. All but one X chromosome are silenced no matter how many are in the cell, nor the sex of the individual [36]. Therefore, only one X chromosome *resists* silencing no matter the number of X chromosomes in the cell.

Clearly, suppressing the *XIST* locus on the future active X is easier for males than females. We know this because of the specific loss of females who reduplicate the essential chromosome 19 gene(s), presumably because reduplication enables both X’s to be active – a known lethal event in diploid cells. At least five percent more preimplantation human females are miscarried than are males [23]. If males reduplicate the *XIST* repressor, it has little consequence, but females who by chance inactivate both *XIST* loci, die before they implant into the uterus. This suggests that not only when this region of chromosome 19 is duplicated, but even, when the chromosome is normal, the required interaction is a difficult one, as either too little or too much *XIST* repressor would lead to a lethal event (too many active X’s or no active X). The former does not occur as often in males who have only one X chromosome: too much repressor is not lethal, although too little might be.

And there is the question of gene dosage. How in a diploid cell do two autosomes cooperate to make an inhibitor for a single X chromosome? And in the case of more than two X chromosomes, how is the right dosage of gene product from chromosome 1 achieved? On one hand, Lyon [37] and more recently Nguyen, Lee & Wu [38] suggest that the two autosomes might pair to synthesize a single product. There is also the possibility of competitive inhibition. Once, a molecule of gene product arrives on one X chromosome then the other(s) are unable to be hit. On the other hand, perhaps, not all attempts to activate or inactivate the chromosome are successful, and so the process is stochastic. That many errors occur while repressing *XIST* on the future active X might explain a significant loss of pre-implantation females, even in absence of gene reduplication.

To answer these questions one needs to identify genome interactions during the pre-implantation development of the human embryo, at the time of X inactivation. One can use chromosome capture such as Hi-C, 3D RNA-FISH [39] (to see if nascent transcripts are transcribed together). And single-cell RNA-Seq as has been recently described in the mouse [26], examining the candidate genes. The best human model would be the beginning of cleavage to embryonic day 10. The inability to study available human embryos is a decided disadvantage for American investigators, but I hope that my colleagues in other countries will carry out such studies. For the human X: 19 interaction, embryonic day 4-7 would probably be appropriate, whereas human embryonic day 6-9 should capture the chromosome 1: X interaction.

## Acknowledgements

The author is most grateful to Drs. Hans Bjornsonn and Teresa Luperchio for incredibly insightful discussions, and to Drs. Haig Kazazian and Roger Reeves for their helpful comments about the manuscript. I am deeply indebted to Dr. Sarah Wheelan for her contribution to the gene conservation analysis.

## Notes

#### Summary of Updates

Table 1 was not clear in the previous submission

